# A TEMPORAL MODEL OF TUMOR-IMMUNE DYNAMICS IN HIGH GRADE SEROUS OVARIAN CANCER

**DOI:** 10.1101/2024.10.21.619446

**Authors:** Donagh Egan, Kate Glennon, Ann Treacy, Aurelie Fabre, Janet McCormack, Salisha Hill, Ryan Morrison, Daniel Liebler, Walter Kolch, Donal Brennan

**Affiliations:** Systems Biology Ireland, University College Dublin, School of Medicine, Belfield, Dublin 4, Ireland; UCD-Gynaecological Oncology Group, School of Medicine, Mater Misericordiae University Hospital, University College Dublin, Dublin, Ireland; Department of Pathology, Mater Misericordiae University Hospital, Dublin, Ireland.; Research Pathology Core Facility, Conway Institute, University College Dublin, Dublin, Ireland; Inotiv, Inc, Nashville, Tennessee; Conway Institute of Biomolecular & Biomedical Research, University College Dublin, Belfield, Dublin 4, Ireland

**Keywords:** Ovarian Cancer, Immune Landscape, Proteomic Profiling, T cell Infiltration, Immunotherapy

## Abstract

Patients with high-grade serous ovarian cancer (HGSOC) typically present with widespread metastasis, obscuring a temporal understanding of tumor-immune dynamics. We performed multi-site global proteomics and matched immunohistochemistry (IHC) of CD4+ and CD8+ tumor infiltrating lymphocytes (TILs) in patient samples. The protein expression profiles were ordered by pseudotime, recapitulating metastatic progression observed in the clinic, and providing a framework to explore tumor-immune dynamics from localized to metastatic disease. Metastatic progression correlated with immune cell infiltration and the recruitment of Tregs to counterbalance the effect of γδ and CD4+ T cells. Whilst later-stage metastases recruited Tregs via chemokines induced by SNX8, early-stage tumors relied more on antigen presentation. The exclusion of CD4+ TILs from the epithelium was correlated with metastatic progression, whereas CD8+ TILs were not, likely due to a predominance of exhausted CD8+ T cells. In early-stage tumors, keratin-expressing cancer cells recruited Tregs via MHC class II, resulting in an inflammatory phenotype characterized by limited IFNγ production and non-clonally expanded T cells. Additionally, IFI44L expression on macrophages caused immune exclusion by downregulating CD53 on T cells. Our findings reveal novel mechanisms of immune escape associated with localized disease and metastatic progression in HGSOC, highlighting potential targets to improve the efficacy of immunotherapy.

## INTRODUCTION

High-grade serous ovarian cancer (HGSOC) typically presents at advanced stages with extensive metastatic spread. This prolonged latency period provides a window for extensive tumor-immune interactions, which are instrumental in shaping both tumor heterogeneity and immunogenicity.^1^ As cancer cells colonize diverse metastatic niches, the tumor-immune relationship evolves, leading to divergent fates among tumors from the same patient.^2^ Despite the clinical significance of this process, a temporal understanding of tumor-immune coevolution from the primary tumor to metastatic disease remains elusive.

Tumor infiltrating lymphocytes (TILs) are a hallmark of immune response and a prognostic marker in HGSOC.^3^ To carry out their function, T cells must directly engage with cancer cells, prompting the use of immunohistochemistry (IHC) to map the spatial relationships between cancer and immune cells. This approach has identified three basic tumor-immune phenotypes: 1) the infiltrated phenotype, characterized by TILs within the tumor epithelium; 2) the immune-excluded phenotype, wherein TILs are restricted to stromal regions surrounding the tumor; and 3) the desert immune phenotype, which lacks TILs entirely.^4^ These immune phenotypes offer a valuable framework for understanding the immune landscape of HGSOC. However, studies have focused on individual tumor sites, leaving the mechanisms that govern the spatial evolution of TILs during metastatic progression largely unexplored.

Conducting longitudinal analyses in clinically relevant HGSOC samples is challenging, as patients often present with tumors at multiple sites, obscuring the temporal sequence of metastatic dissemination. However, pseudotime analysis can approximate temporal variation by projecting samples into a unidimensional latent space that quantifies relative disease progression.^5^ Normally used in single-cell RNA-seq, we have adopted this approach for analyzing multi-site biopsies commonly obtained from surgical debulking of HGSOC.

To investigate tumor-immune crosstalk during metastatic progression, we performed multi-site global proteomic profiling on 80 tumor samples collected from 23 patients (Figures 1A and 1B). The spatial distribution of CD4+ and CD8+ TILs was mapped using IHC in a subset of 21 samples from 7 patients, enabling us to correlate molecular changes with the spatial dynamics of TILs. Additionally, we assessed intra-tumor heterogeneity (ITH) by expanding the number of regions analyzed by IHC in primary ovarian and omental tumors from 5 patients (n = 10 samples), generating a detailed map of immune phenotypes across different tumor regions.

**Figure 1.**
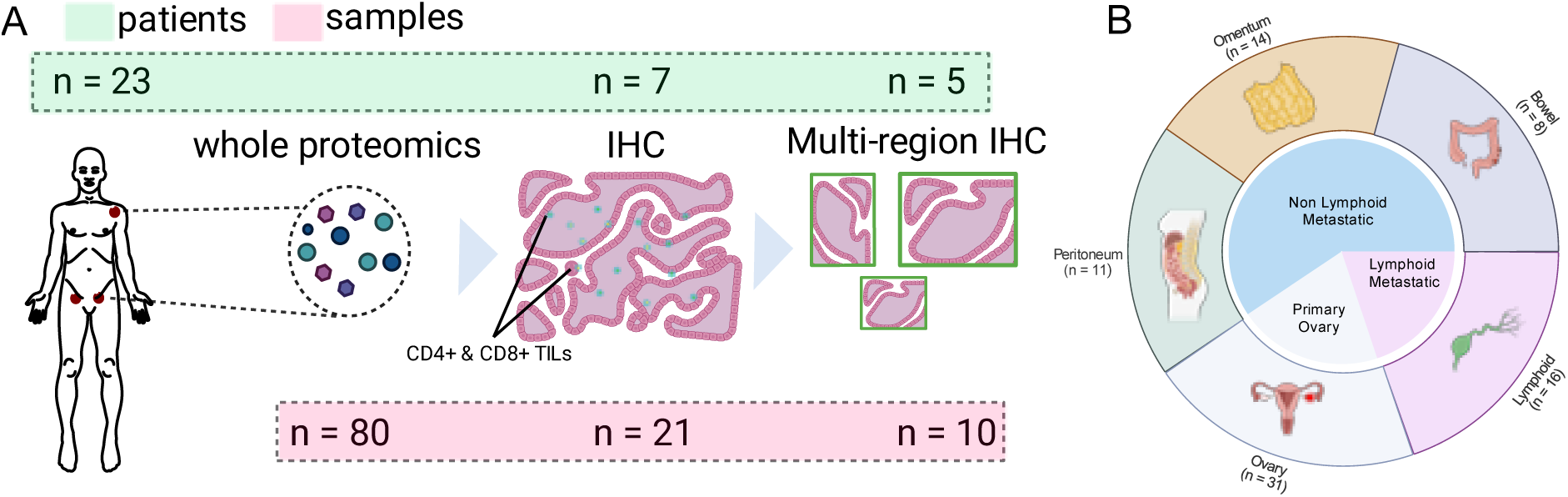
Patient Sampling and Study Design. A) The methods used for profiling tumors, and number of patients (top box) and samples (bottom box) available for each. B) The number of samples for global proteomic profiling, and their broader classification to either primary ovarian, lymphoid metastases, and non-lymphoid metastases.

## RESULTS

### The Recruitment of Tregs Drives Metastatic Progression

We conducted global proteomics on Formalin-Fixed Paraffin-Embedded tumor sections (n = 23 patients/80 samples) (Supplemental Table S1). In total, 5,824 unique proteins were identified (median/sample = 4,654), aligning with the coverage reported in prior studies.^6^ Furthermore, the number of proteins identified and their dynamic ranges were consistent across tumor sites (Supplementary Figure 1A and 1B).

Principal component analysis (PCA) identified metastasis status and tumor site as the primary sources of variation, rather than patient identity (Figure 2A) (Supplementary 1C). To create a framework for understanding metastatic progression, we performed pseudotime analysis with Phenopath,^5^ projecting each sample into a continuous, patient-specific latent space (Figure 2B). The pseudotime scores increased from primary to metastatic samples (p < 0.05) (Figure 2C), with tumor sites positioned in line with clinical observations of metastatic spread: primary ovarian samples first, followed by the omentum and peritoneum due to direct dissemination within the abdominal cavity, then the bowel via peritoneal spread, and finally the lymph nodes (Figure 2D).^7^

**Figure 2.**
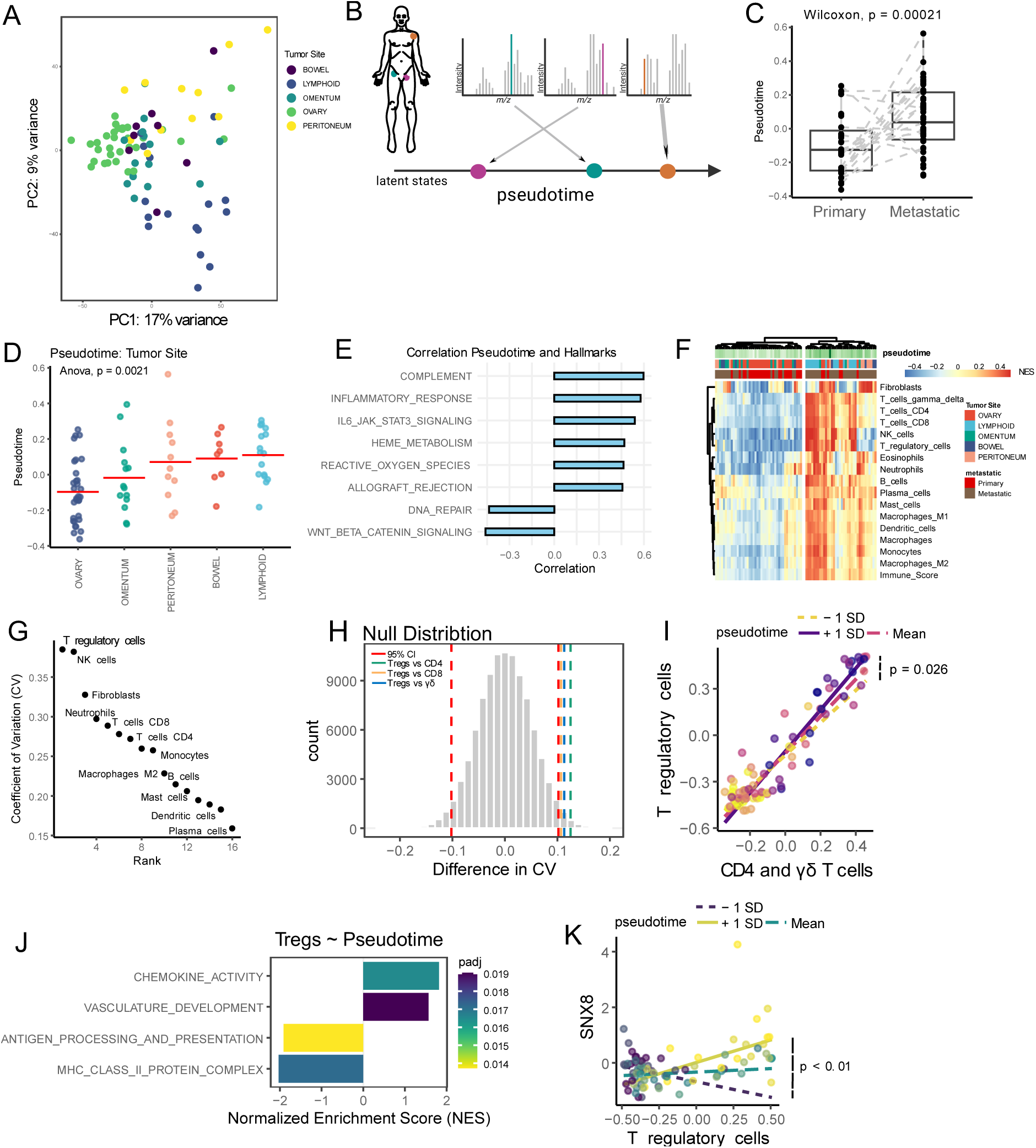
Pseudotime Modeling of Metastatic Progression in HGSOC. **A)** Principal component analysis on the proteomic profiles of HGSOC tumor samples, annotated by tumor site. **B)** A schematic of the pseudotime analysis in multiple samples from one patient to identify temporal information that traces metastatic progression in HGSOC from a cross-sectional cohort. **C)** Pseudotime estimates in primary and metastatic samples. **D)** Pseudotime estimates in samples from distinct tumor sites. **E)** Spearman’s rho between the pseudotime estimates and pathway activity scores inferred using single sample GSEA. The hallmark pathways were used, and the top 8 most significant pathways were selected. **F)** Hierarchical clustering of HGSOC tumor samples according to deconvolution derived estimates of different cell types. The abundance of each cell type is represented by a normalized enrichment score (NES). **G)** The coefficient of variation (CV) for each cell type across the tumor samples. **H)** A null distribution representing the differences in CV between cell types, estimated using permutations (n = 100,000) without replacement. The red lines represent the 95% confidence interval; the remaining lines represent comparisons between cell types. **I)** The correlation between the mean abundance of effector CD4+ T cells and γδ T cells with Tregs, assessed across samples with different pseudotime estimates (-1 standard deviation; mean; and +1 standard deviation) **J)** Enriched pathways for genes whose correlation with Tregs changes as a function of pseudotime. A positive NES indicates pathways where the correlation with Tregs increases with pseudotime, while a negative NES reflects pathways where this correlation decreases. **K)** The correlation between SNX8 expression and Treg abundance for samples with different pseudotime estimates (-1 standard deviation; mean; and +1 standard deviation).

To identify pathways involved in metastatic progression, we calculated Spearman’s rho between the pseudotime estimates and pathway activity scores (Hallmarks database), inferred using single sample GSEA. Immune related pathways such as the complement system, inflammatory response, and IL6 signaling trough JAK and STAT, were the most significant pathways positively correlated with pseudotime (Figure 2E) (FDR < 0.05). IL-6 produced by omental adipocytes promotes invasion of ovarian cancer cells,^8^ whereas the complement system and inflammatory response reflect escalating immune dysfunction, wherein cancer cells hijack macrophages and Tregs to create an immunosuppressive environment.^9^

We explored the relationship between tumor composition and metastatic progression by estimating the abundance of various cell types using a deconvolution algorithm.^10^ Hierarchical clustering grouped the samples based on tumor site, metastatic status, and pseudotime, revealing increased immune cell infiltration at later pseudotime scores (Figure 2F). Tregs emerged as the most variable cell type across the samples, exhibiting significantly greater variability compared to effector T cells (p < 0.05) (Figure 2G and 2H). We hypothesized that this variability in Tregs may reflect their migration with effector T cells, serving to mitigate immune predation at metastatic sites. Indeed, as pseudotime progressed, the correlation of Tregs with CD4+ and γδ T cells increased significantly (p = 0.026) (Figure 2I). However, this was not observed for CD8+ T cells, suggesting that Treg recruitment to metastatic sites is particularly crucial for mitigating CD4+ and γδ T cell responses.

To investigate the mechanisms underlying Treg recruitment during metastatic progression, we identified genes whose correlation with Tregs changed as a function of pseudotime, and mapped these to functional pathways. As pseudotime advanced, we observed a significant increase in the correlation between Tregs and pathways involved in chemokine signaling and vasculature development, but a decrease in the correlation with antigen presentation pathways (FDR < 0.05) (Figure 2J). This indicates that early-stage tumors predominantly accumulate Tregs through antigen presentation, while metastatic sites rely on chemokine-mediated signaling and vascularization.

From this analysis, the correlation between SNX8 and Tregs increased the most with pseudotime (FDR = 0.003) (Figure 1K, Supplementary Figure 1D). SNX8 switches IFNγ signaling to the noncanonical pathway, upregulating the expression CXCL9 and CXCL10, chemokines that recruit Tregs through CXCR3.^11,12^ We confirmed the coregulation of SNX8 with Tregs, CXCL9/10, and CXCR3 in ovarian cancer samples from The Cancer Genome Atlas (TCGA) (Supplementary Figure 1E). Furthermore, the correlation between IFNγ signaling and the expression of Treg-recruiting chemokines was stronger in tumors with high SNX8 expression (Supplementary Figure 1F). These findings suggest that immune-infiltrated tumors leverage the IFNγ-SNX8-CXCL9/10 axis to recruit Tregs, creating an infiltrated, but immunosuppressive and inflammatory environment.

### CD4+ TILs Are Excluded from the Epithelium During Metastatic Progression

We mapped the spatial distribution of CD4+ and CD8+ TILs in intraepithelial (iTILs) and stromal (sTILs) regions using IHC (n = 7 patients/23 samples) (Supplemental Table S2). A positive correlation was observed between the density of iTILs and sTILs for both CD4+ (R = 0.83; p < 0.01) and CD8+ (R = 0.72; p < 0.01) TILs (Figure 3A and 3B). Therefore, the density of TILs that infiltrate the epithelium is strongly related to the densities observed in the stroma.

**Figure 3.**
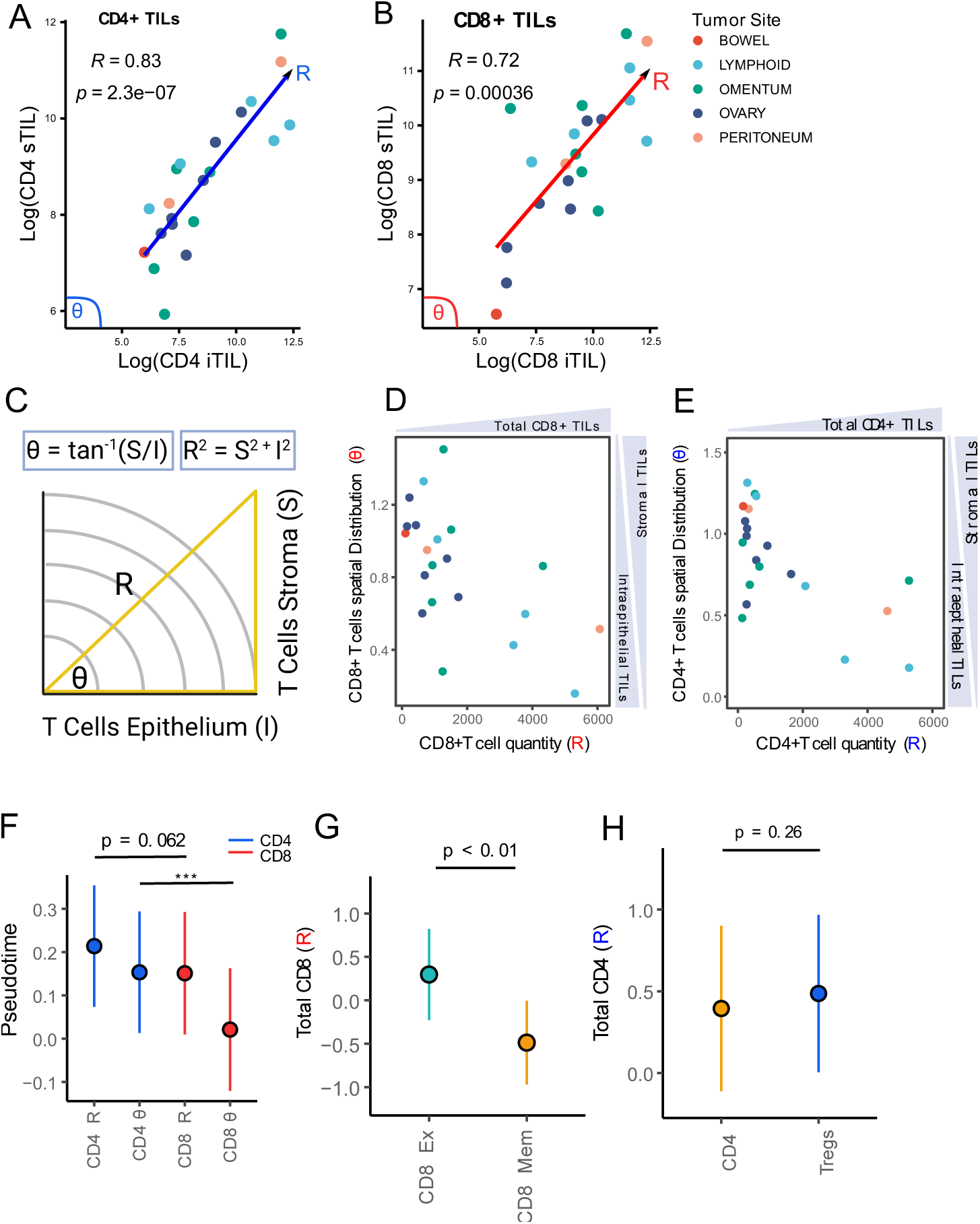
The Spatial Distribution of TILs During Metastatic Progression. **A, B)** Spearman’s rho between intraepithelial TILs (iTILs) and stromal TILs (sTILs) for CD4+ and CD8+ T cells, respectively. **C)** The method for calculating the polar coordinates. The R value quantifies the overall density of iTILs and sTILs, defined as R = square root [(TILs tumor)2 + (TILs stroma)2]. θ quantifies the spatial distribution of TILs in the stoma relative to the epithelium, defined as θ = atan(TILs stroma/TILs tumor). **D, E)** The overall density (R) and spatial distribution (θ) of CD4+ and CD8+ TILs, respectively. **F)** Coefficient estimates for the association of the polar coordinates with pseudotime, compared using a t-test. **G, H)** Coefficient estimates for the association of CD8+ and CD4+ T cells phenotypes with the overall density of CD8+ and CD4+ TILs (R), respectively. Significance was determined using a t-test. The vertical lines represent the 5% and 95% confidence intervals (CIs). P-values from the multiple regression (*** = p < 0.01).

To better understand the spatial distribution of TILs we created a new polar coordinate system. Briefly, the densities of CD4+ and CD8+ TILs were converted into two new metrics: 1) The overall density of CD8+ and CD4+ T cells (R value), and 2) the distribution of CD8+ and CD4+ T cells in the epithelium compared to stromal compartments (θ value) (detailed in methods; adopted from^13^) (Figure 3C-3E).

We explored the spatial evolution of TILs during metastatic progression by analyzing the relationship between pseudotime and the polar coordinates. Consistent with our previous findings (Figure 2F), the overall density of CD8+ and CD4+ TILs (R value) correlated with pseudotime (p = 0.012 and p = 0.004, respectively). However, the overall abundance of CD4+ TILs (R value), and their exclusion from the epithelium (θ value), demonstrated a stronger association with metastatic progression compared to CD8+ TILs (p = 0.062 and p < 0.01, respectively) (Figure 3F).

Using gene signatures from a published study,^14^ we found that CD8+ TILs consisted of significantly more exhausted CD8+ T cells compared to memory T cells (p < 0.01) (Figure 3G). Given that memory T cells are essential for anti-tumor immunity, and exhausted T cells have limited functionally capacity,^14^ their exclusion from the epithelium likely has minimal impact on metastatic progression. In contrast, Tregs and CD4+ effector T cells contributed equally to the overall density of CD4+ TILs (Figure 3H). Tregs were also significantly correlated with the exclusion of CD4+ TILs from the epithelium (θ value, p = 0.03), suggesting they play a key role in driving this process and its link to metastatic progression.

### CD4+ and CD8+ TILs Exhibit Distinct Patterns of Heterogeneity

In HGSOC, TILs exhibit greater intratumor heterogeneity (ITH) than intra-patient heterogeneity, posing an obstacle for immunotherapy treatments.^15^ We investigated ITH in early-stage primary and omental tumors by analyzing each IHC slide at high magnification (x20), and quantifying the number regions classified as infiltrated, excluded, or desert (Figure 4A) (Supplemental Table S3). For example, at a low magnification, a full-face section was initially divided into five major regions and classified as excluded due to a higher density of sTILs compared to iTILs. However, when the magnification was increased, 26 individual regions were identified, consisting of infiltrated (n = 12), excluded (n = 6), and desert (n = 16) immune phenotypes (Supplementary Figure 2A-E).

**Figure 4.**
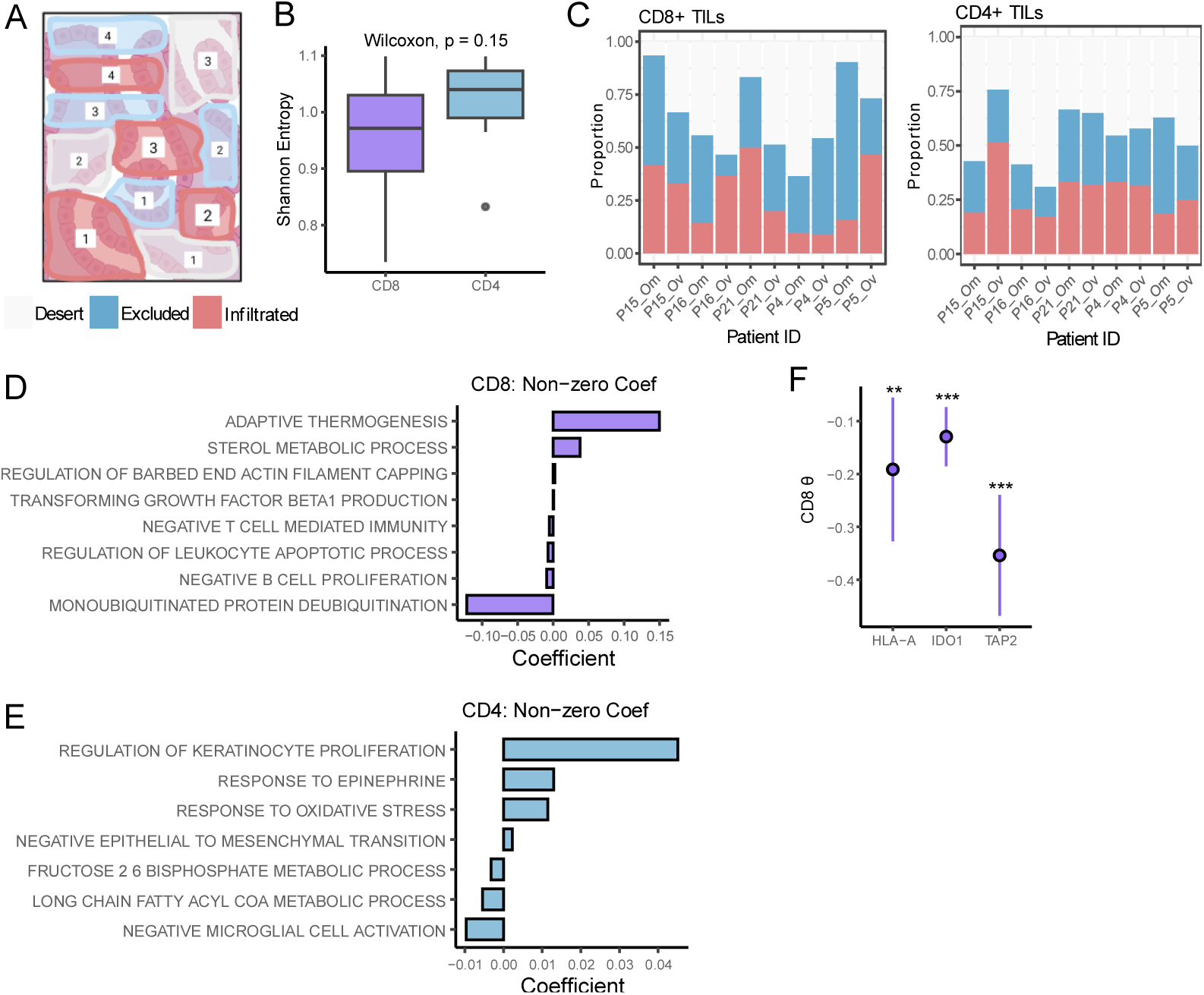
Intratumor Heterogeneity of The Immune Phenotypes. **A)** An illustrative IHC slide, analyzed at high magnification (x20), where the number of immune, excluded, and infiltrated regions are quantified. **B)** The Shannon diversity index for the abundance of immune phenotypes in a tumor, assessed for both CD8+ and CD4+ TILs**. C)** The distribution of immune phenotypes for CD8+ and CD4+ TILs, respectively. The association between the immune phenotype distributions and tumor samples was assessed using a chi-squared test. **D, E)** The non-zero coefficients from an elastic net regression, obtained by modeling pathway activity scores against the SDI for CD8+ and CD4+ TILs, respectively. **F)** Coefficient estimates for the association of HLA-A, IDO1, and TAP2 with CD8+ TIL infiltration (θ). The vertical lines represent the 5% and 95% CIs.

We used the Shannon diversity index (SDI) to formally quantify the ITH of immune phenotypes in each sample. In this context, a lower SDI reflects a tumor dominated by a single immune phenotype, while a higher SDI signifies a more balanced distribution of immune phenotypes. CD8+ TILs trended toward a lower SDI compared to CD4+ TILs (p = 0.14) (Figure 4B). Furthermore, the distribution of immune phenotypes showed greater variability across tumor samples for CD8+ TILs compared to CD4+ TILs (p < 0.01 and p = 0.06, respectively) (Figure 4C).

We used Elastic Net regression to identify the pathways that best explain the variation in SDI values. After regularization, leukocyte apoptosis, negative regulation of T cells, and TGBF1 production remained as independent predictors for CD8+ TILs (Figure 4D). In contrast, oxidative stress and keratinocyte proliferation were selected for CD4+ TILs (Figure 4E). Therefore, we hypothesized that antigen engagement, coupled with co-inhibitory signals, may underlie the low ITH observed for CD8+ TILs. To test this, we analyzed a dataset published by Zhang et al.,^1^ which mapped the spatial distribution of CD4+ and CD8+ TILs in the epithelium and stroma of HGSOC patients (n = 109). Indeed, CD8+ TIL infiltration (θ value, as previously defined) was significantly correlated with the expression of antigen-presentation genes, including TAP2 and HLA-A, as well as the endogenous immune checkpoint molecule IDO1 (p < 0.05) (Figure 4F). Collectively, these findings demonstrate that the ITH, and the pathways regulating it, are distinct between CD4+ and CD8+ TILs.

### Keratin Expressing Cancer Cells Constrain the Immune Response Via Tregs

Having identified mechanisms that drive metastatic progression and ITH in early-stage disease, we analyzed the omental and ovarian cancer samples to elucidate novel tumor-immune interactions that permit localized disease. The densities of CD8+ and CD4+ TILs, determined by IHC analysis, were significantly correlated with deconvolution-derived estimates of CD8+ and CD4+ T cells in the global proteomics dataset (p < 0.05) (Supplementary Figure 3A), highlighting a concordance and mutual robustness of both approaches. Therefore, we categorized each sample as infiltrated or non-infiltrated based on the predominant immune phenotype from the multi-region IHC analysis, and identified differentially expressed proteins between both groups.

Keratin proteins (KRT 1, 2, 9, and 10) were the most upregulated proteins in infiltrated tumors, whereas IFI44L was the most upregulated protein in non-infiltrated tumors (Figure 5A). As expected, immune-related pathways, such as adaptive immune response and lymphocyte activation were enriched in infiltrated tumors. Conversely, the spliceosome pathway was downregulated, potentially enhancing tumor immunogenicity by creating more splicing-derived neoantigens (Figure 5B).^16^

**Figure 5.**
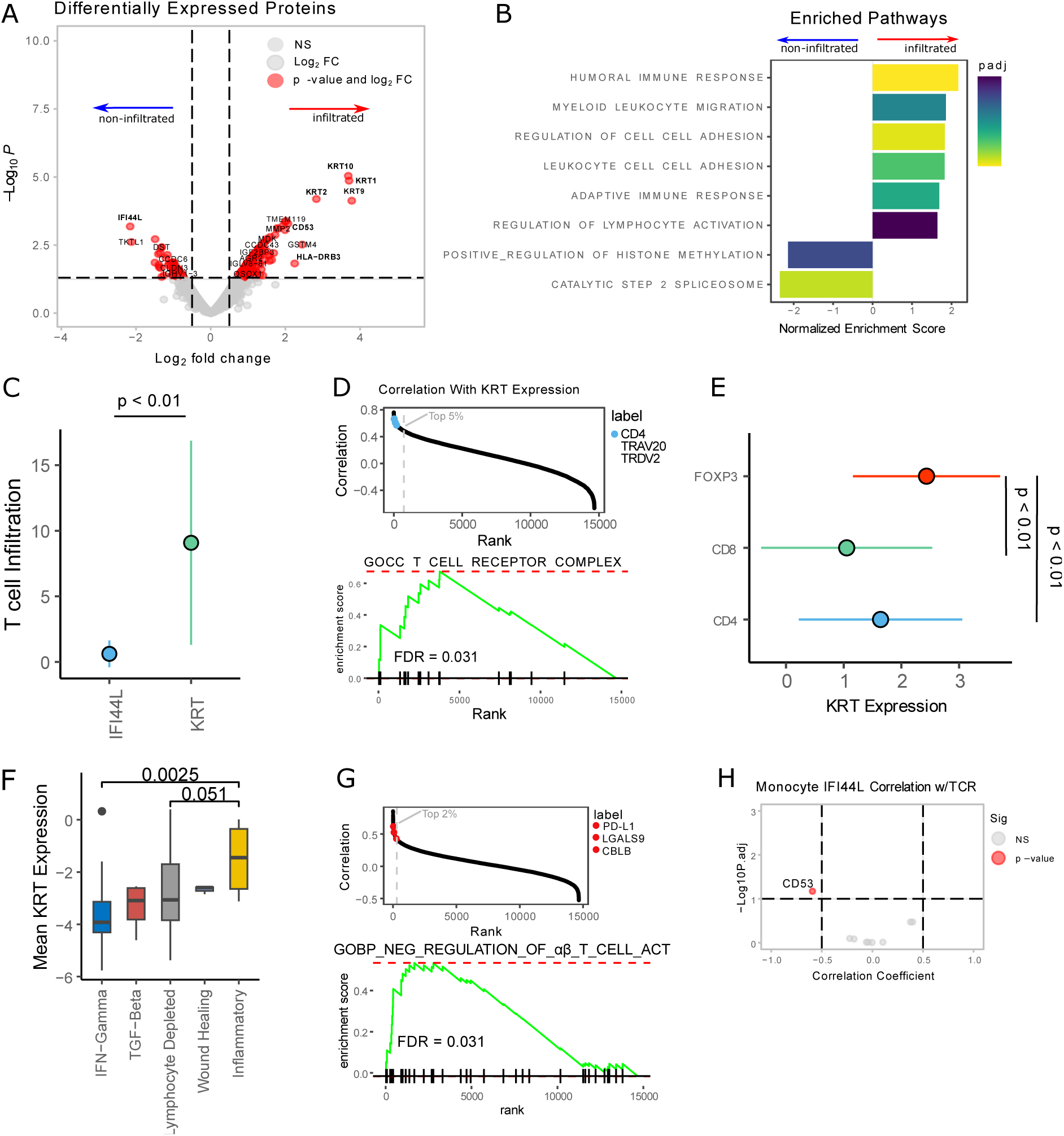
The Mechanisms Underlying T Cell Infiltration and Exclusion. **A)** The differentially expressed proteins between infiltrated and non-infiltrated tumors. The genes delineated in red satisfied the significance (p value < 0.05) and fold change (log2FC > 1) thresholds. **B)** Enriched pathways between infiltrated and non-infiltrated tumors. An FDR threshold of 0.05 was implemented. **C)** Coefficient estimates for the association of keratin and IFI44L expression with the fraction of T cells in the published dataset,^15^ compared using a t-test. The vertical lines represent the 5% and 95% CIs. **D)** Pearson’s correlation coefficient between each gene and keratin expression (top panel). Genes left of the dotted gray line are in the top 5%. GSEA for the T cell receptor complex pathway using genes ranked by their correlation with keratin expression (bottom panel). **E)** Coefficient estimates for the association between keratin expression and the fractions of Tregs, CD4+, and CD8+ T cells in the published dataset,^15^ compared using a t-test. The horizontal lines represent the 5% and 95% CIs. **F)** The mean keratin protein expression in the independent proteogenomic ovarian cancer dataset.^21^ The tumor phenotypes are compared using a Wilcoxon rank sum test. **G)** Pearson’s correlation coefficient between each gene and IFI44L expression (top panel). Genes left of the dotted gray line are in the top 2%. GSEA for the negative regulation of αβ T cell activation pathway using genes ranked by their correlation with IFI44L expression (bottom panel). **H)** The correlation between receptor expression in T cells and IFI44L expression in Monocytes/Macrophages. The gene delineated in red satisfied the significance (FDR < 0.05) and correlation coefficient (R > |0.5|) thresholds.

As an initial validation step, we confirmed the expression of keratin proteins in ovarian cancer cells using two independent single-cell RNA-seq datasets, which were integrated with Harmony (Supplementary Figure 3B-3D).^17–19^ We then leveraged a published dataset with multicolor immunofluorescence quantification of CD4+, CD8+, and Tregs in treatment-naïve HGSOC samples (n = 37).^15^ At least ten tumor sections per sample were analyzed, excluding stromal areas, yielding 440 regions with matched transcriptomic data. In this dataset, we confirmed that keratin expression (KRT1, KRT2, KRT9, and KRT10) is correlated with T cell infiltration (p = 0.02), and this association is significantly stronger compared to IFI44L (p < 0.01) (Figure 5C).

To understand keratin-induced T cell infiltration, we identified co-expressed genes using Pearson’s correlation in the validation dataset and mapped them to Gene Ontology pathways with GSEA. The co-expressed genes were significantly enriched for the T cell receptor complex pathway, with CD4, TRAV20, and TRDV2 among the top five percent of co-expressed genes (FDR < 0.05) (Figure 5D). Keratin expressing stromal cells constrain the immune response by presenting self-antigens via MHC class II to Tregs.^20^ Given the co-expression of keratin proteins with variable TCR regions and the concomitant upregulation of HLA-DRB3 in infiltrated tumors, we hypothesized that ovarian cancer cells retain this mechanism from their epithelial origin (Figure 5A). Indeed, keratin expression was the strongest predictor of Treg infiltration, followed by CD4+ T cells, and then CD8+ T cells in (p < 0.01) (Figure 5E).

Next, we analyzed keratin protein expression in a proteogenomic dataset of ovarian cancer samples (n = 82) classified according to phenotypes from the TCGA pan-cancer study.^21,22^ Tumors with an inflammatory phenotype upregulated keratin expression compared to lymphocyte-depleted and IFNγ-expressing tumors (p = .051 and p = .0025, respectively) (Figure 5F). This demonstrates that keratin expression is associated with inflammation, infiltrating lymphocytes, but not with interferon gamma production, supporting its role in constraining the immune response through Tregs. Finally, high keratin expression was associated with worse overall survival in ovarian cancer datasets from TCGA and gene expression omnibus (GEO) (p < 0.05) (Supplementary Figure 3E), demonstrating that Treg infiltration due to keratin expression is associated with worse prognosis in HGSOC.^23^

IFI44L is an interferon-stimulated gene that promotes macrophage differentiation and downregulates the immune response following viral and bacterial infections.^24,25^ Its expression was not associated with T cell infiltration in the validation dataset,^15^ but was co-expressed with genes involved in negative T cell regulation, including PD-L1, LGALS9, and CBLB (FDR < 0.05) (Figure 5G). Furthermore, its expression was strongly upregulated in Monocytes/Macrophages in the previously described single cell RNA-seq datasets (p < 0.05) (Supplementary Figure 3F). To investigate an IFI44L-mediated functional relationship between Monocytes/Macrophages and T cells, we correlated IFI44L expression in Monocytes/Macrophages with receptor expression in T cells, as defined by Omnipath.^26^ A negative correlation between CD53 expression in T cells and IFI44L expression in Monocytes/Macrophages was observed (FDR < 0.05) (Figure 5H). Furthermore, CD53 expression was downregulated in non-infiltrated samples, consistent with an inverse relationship with IFI44L (Figure 5A). These findings suggest that IFI44L inhibits CD53 expression in T cells. Since CD53 promotes T cell extravasation,^27^ its downregulation likely contributes to immune exclusion. Finally, IFI44L expression trended with reduced overall survival in TCGA and GEO datasets (p = 0.069) (Supplementary Figure 3G).^23^

### The Proteome Underlying Keratin Expression Restricts T Cell Clonal Expansion

Upon antigen-specific T-cell receptor stimulation, Tregs suppress the cytotoxic activity of CD8+ T cells and restrict clonal expansion against both self and foreign antigens.^28^ To investigate how the proteome underlying elevated keratin expression impacts the clonal composition of immune cells, we re-analyzed the Zhang et al. dataset,^1^ leveraging the matched transcriptomic, T- and B-cell receptor sequencing data from 111 samples across 20 HGSOC patients. These samples were previously classified into distinct immune phenotypes: desert, excluded, and infiltrated.

First, we validated that the proteins identified as upregulated (logFC > 0, p < 0.05) in the infiltrated samples from our dataset, referred to as the “infiltrated signature,” were significantly correlated with the immune-infiltrated phenotype in the published dataset.^1^ (p < 0.05) (Figure 5A). Furthermore, the proteins identified as upregulated in non-infiltrated samples (logFC < 0, p < 0.05) - termed the “non-infiltrated signature” – were correlated with the desert immune phenotype (p = 0.06) (Figure 6A). These findings suggest that the proteomic changes underlying elevated keratin expression are conserved at the transcriptomic across independent datasets.

**Figure 6.**
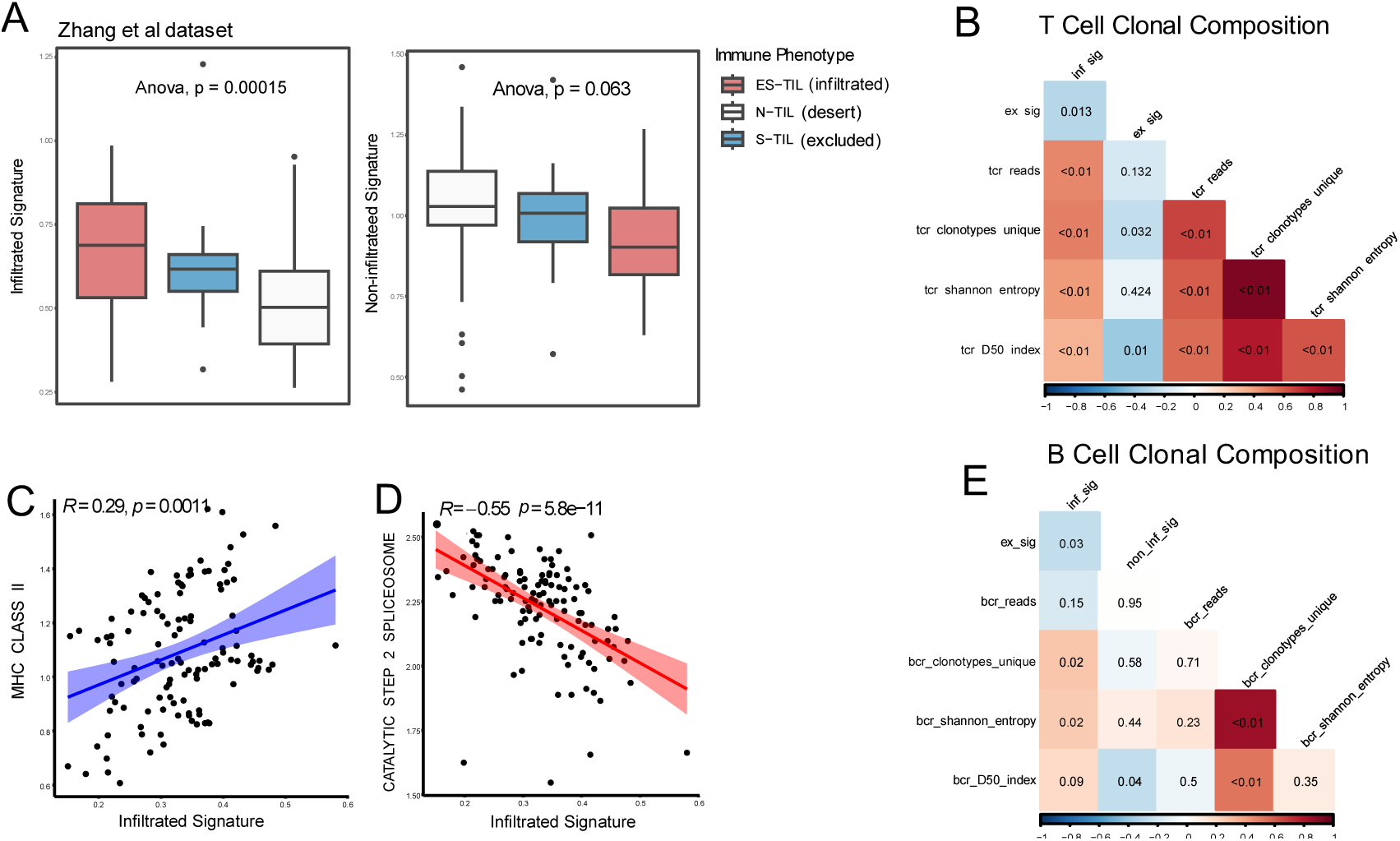
The Proteome Underlying Keratin Expression Restricts Clonal Expansion. **A)** The activity of proteins upregulated in infiltrated tumors (left panel) and non-infiltrated tumors (right panel) in the published dataset. The samples were classified by the original authors using IHC. **B)** The correlation of the infiltrated signature (inf_sig) with T cell abundance (tcr_reads), T cell diversity (Shannon entropy, D50 index), and the number of unique T cell clones (clonotype unique). The color scale represents the correlation coefficients and the p values are provided in the tiles. **C, D)** The correlation between the infiltrated signature and the activity of the MHC class II pathway and the catalytic spliceosome pathway, respectively. **E)** The correlation of the infiltrated signature (inf_sig) with B cell abundance (bcr_reads), B cell diversity (Shannon entropy, D50 index), and the number of unique B cell clones (clonotype unique). The color scale represents the correlation coefficients, and the p values are provided in the tiles.

Next, we analyzed T and B cell receptor sequencing data. Given that individual patient contributed multiple samples, we calculated a partial Spearman’s correlation coefficient between the previously defined infiltrated and non-infiltrated signatures and metrics defining the clonal composition of T and B cells. The infiltrated signature positively correlated with the number of TCR reads in a sample, a proxy for T cell infiltration, as well as measures of clonal diversity such as Shannon’s entropy and the D50 index (p < 0.05) (Figure 6B). This suggest that keratin proteins promote T cell infiltration, but restrict clonal expansion (emergence of dominant clone), indicative of Treg-induced immunosuppression.

In our previous analysis, we proposed two mechanisms underlying T cell infiltration in infiltrated samples: 1) increased splicing-derived neoantigens due to spliceosome downregulation, and 2) MHC class II antigen presentation by keratin-expressing cells to Tregs. In the published dataset,^1^ we confirmed a positive correlation between the infiltrated signature and MHC class II expression, and a negative correlation with spliceosome catalytic activity (Figure 6C and 6D).

Tregs are essential and sufficient to suppress autoreactive B cells in an antigen-specific manner and to prevent them from producing autoantibodies.^29^ The infiltrated signature demonstrated no correlation with B cell infiltration (BCR reads) (p > 0.05), but was positively correlated with measures of clonal diversity (p < 0.05) (Figure 6E). Therefore, although the proteome underlying keratin expression does not promote B cell infiltration, it is capable of restricting their clonal expansion, likely through the action of Tregs.

## DISCUSSION

Immune checkpoint blockade has failed to deliver meaningful responses in HGSOC, despite being considered an immunogenic disease.^30,31^ To improve patient outcomes, a temporal understanding of tumor-immune coevolution is required. We addressed this by projecting multi-site HGSOC samples into a pseudotime latent space that recapitulates clinical observations of metastatic dissemination. Within this novel framework, we identified mechanisms of immune escape from a localized to metastatic phenotype.

Tregs are known to foster an immune privilege that promotes tumor progression.^32–35^ However, their specific role in the pathogenesis and metastatic progression of HGSOC remains understudied due to limited multi-site datasets. We demonstrated that immune cell infiltration increases with metastatic progression, accompanied by the recruitment of Tregs to counteract CD4+ and γδ T cells. For CD4+ TILs, Tregs were implicated in their exclusion from the epithelium. As γδ T cells display a cytotoxic phenotype without expressing immune checkpoints such as PD1 and CTLA4,^36^ metastatic tumors may rely on Tregs to suppress their cytotoxic activity.

Tregs accumulate in ovarian cancer tumors either by recognizing neoantigens and undergoing clonal expansion,^37^ or through chemokine-mediated recruitment.^35^ Our results demonstrate that early-stage tumors rely more on antigen presentation to recruit Tregs, whereas later-stage metastases increasingly depend on chemokine signaling and vasculature development. We posit that the frequent loss of HLA expression observed in metastasizing ovarian cancer cells drives a shift away from antigen-driven Treg recruitment.^38^

SNX8 acts as a molecular switch in noncanonical IFNγ signaling, promoting the upregulation of key chemoattractants for Tregs, such as CXCL9 and CXCL10.^11,12^ At metastatic sites, where immune infiltration and IFNγ levels are elevated, this mechanism may be exploited by cancer cells to enhance Treg recruitment, thereby facilitating immune suppression. Furthermore, IFNγ induced by immune checkpoint blockade may inadvertently promote Treg recruitment through this pathway.

Unlike CD4+ TILs, the exclusion of CD8+ TILs from the epithelium was not associated with metastatic progression, likely reflecting their predominately exhausted phenotype. Our findings in early-stage tumors highlighted antigen presentation and immune inhibitory molecules as key factors shaping the spatial distribution and ITH of CD8+ TILs. Therefore, driving CD8+ T cell exhaustion through antigen exposure and immune checkpoint activation, rather than physical exclusion, may be the primary mechanism diminishing their cytotoxic potential. Since immune checkpoint blockade has demonstrated limited capacity for reinvigorating exhausted T cells, instead promoting the infiltration of novel T cell clones,^39,40^ it will be important to investigate the migratory capacity of the T cell repertoire from the periphery in HGSOC.

Single cell RNA-seq of the fallopian epithelium identified a cell phenotype characterized by keratin expression and upregulation of MHC class II genes.^41^ In a mouse model, keratin expressing stromal cells present antigens via MHC class II, maintaining an active population of Tregs that constrains antigen-specific immunity.^20^ We demonstrated that ovarian cancer cells in early-stage disease likely retain this mechanism from their epithelial origin. This results in an inflammatory phenotype characterized by limited IFNγ production, restricted B and T cell clonal expansion, and ultimately worse patient prognosis.

IFI44L’s function has primarily been described in the context of viral and bacterial infections, where it suppresses innate immune responses triggered by viral infections.^24,25^ Therefore, it has been proposed to treat diseases with excessive IFN production. We propose a novel function of IFI44L in promoting immune exclusion in HGSOC through an inhibitory relationship with CD53 expression in T cells. Given its co-expression with PD-L1, a primary target of immune checkpoint blockade, IFI44L could serve as a novel target for future immune checkpoint blockade therapies.

Overall, this study identifies mechanisms of immune evasion that permit localized disease and drive metastatic progression in HGSOC. The aberrant role of Tregs at each stage of disease underscores the importance of directly targeting this cell type to enhance immunotherapy outcomes for HGSOC patients.

## MATERIAL AND METHODS

### Immunohistochemistry

The tissue was collected during laparoscopic biopsy or primary cytoreductive surgery, then formalin-fixed and paraffin-embedded. Sections were stained with hematoxylin and eosin, visualized with diaminobenzidine, and counterstained with hematoxylin per the manufacturer’s instructions. The slides were dehydrated, cleared, and permanently mounted. This involved 4μm thick tissue sections mounted onto a “charged “glass slide and baked at 60 degrees for 1 hour. The slides were then labelled and loaded on the Leica Bond III immunostainer. The sections were stained with either CD4 (Leica Ready to Use antibody CD4 PA0427 Clone 4B12) or CD8 (Leica Ready to Use antibody CD8 PA0183 Clone 4B11) on an IHC autostainer. The slides were dewaxed, antigens were retrieved, and then the slides were dehydrated, cleared, and coverslipped. The stained slides scanned using an Aperio AT2 digital slide scanner (Leica Biosystems Ltd, Germany). The scanned images were annotated with a consultant pathologist to assess intra-epithelial and stromal CD4 and CD8 expression using ImageScope (Leica Biosystems Ltd), and analyzed using the Aperio Nuclear Algorithm v9 (Leica Biosystems Ltd). Ovarian tumor epithelial cells were distinguished from stromal cells by the shape and size of their nuclei. The TIL density was calculated as the number of CD4 and CD8 positive cells per mm2.

### Global Proteomics Analyses

High resolution liquid chromatography-tandem mass spectrometry (LC-MS/MS) analysis with Tandem Mass Tag (TMT) labeling was used, as described previously with modifications.6 FFPE tissues were deparaffinized and proteins were extracted and digested with trypsin as described.7 Two micrograms of protein from each sample were combined to produce a common control reference sample and 20 μg of protein from each sample and a 20 μg aliquot of the common control were TMT labeled with ThermoFisher Scientific 18-plex reagent kits according to the manufacturer’s instructions. Samples were combined in equal amounts and separated into 8 fractions with high pH reverse phase fractionation spin columns. LC-MS/MS analyses of each fraction were performed with an Orbitrap Fusion Lumos mass spectrometer coupled to an Ultimate 3000 nano-LC. Peptides were separated over a 3 hr gradient from 4 to 25% acetonitrile in 0.1% formic acid at a flow rate of 300 nL/min on a Thermo Scientific EasySpray 50 cm x 75 µm, 2 µm particle size RP C18 column. Full scan MS spectra were acquired in the Orbitrap at a resolution of 120,000. The most intense MS1 ions were selected fragmentation in the ion trap at 35% collision energy and an isolation width of 1.2 Da. Six fragment ions were co-isolated using synchronous precursor selection with an isolation width of 1.3 Da and fragmented by HCD with a normalized collision energy of 65%. The product ions were analyzed using the Orbitrap at a resolution of 50,000. Peptide and protein identifications and relative quantification were done with Proteome Discoverer® v. 2.5 software.

The relative abundances were log2 transformed and zero-centered for each protein to obtain final, relative abundance values. Proteins absent in 50% or more of samples in any condition were filtered out, followed by imputation using the “scImpute” function and tail-based imputation using the “tImpute” function from PhosR.

### Pseudotime Analysis

Pseudotime analysis was performed on global proteomic data from 23 patients (n=80 samples) encompassing both primary and metastatic tumors, using the PhenoPath R package.^5^ To account for multiple samples per patient, patient identity was included as a covariate in the model. Pathway associations were assessed by calculating Spearman’s rho between pseudotime and hallmark pathway activity scores (inferred with single-sample GSEA). Benjamini-Hochberg correction was applied to adjust for multiple testing.

### TME Deconvolution

TME cell composition was estimated using the ConsensusTME method with ovarian cancer-specific gene sets.^10^ The coefficient of variation for each cell type was calculated. To assess the significance of cell type variation differences, a null distribution was generated through a permutation test (100,000 replicates) without replacement.

### The Polar Coordinate System

The density of CD4+ and CD8+ TILs in the epithelium and stroma were converted into polar coordinates, defined by the following equations: 1) T cell quantity (R) = [square root ((TILs tumor)2 + (TILs stroma)2)]; 2) T cell spatial distribution (θ) = [atan (TILs stroma/TILs tumor)]. Multiple linear regression was used to determine the coefficients between the polar coordinates, pseudotime and T cell phenotypes, which were then compared using a T-test. The signatures for CD8+ T cell phenotypes were accessed from the supplemental data of Sade Feldman et al,^14^ and were scored in each sample using single sample GSEA.

### Shannon Diversity Index Analysis

The number of desert, excluded, and infiltrated regions for CD4+ and CD8+ TILs were quantified in each sample. These counts served as input for the entropy.empirical function in the ‘entropy’ R package, employing the maximum likelihood estimation method with default parameters. Pathways from the ontology gene set, available through MSigDB, were scored in each sample, and their relationship with the SDI values was modeled using elastic net regression via the glmnet function in R. The lambda that minimized the cross-validated error was selected, with an alpha value of 0.5 to balance both Lasso and Ridge regression penalties. Only non-zero coefficients were retained for further analysis.

### Single cell RNA-seq Integration and Cell-cell Communication Analysis

Two independent single cell RNA-seq datasets were accessed from the TISCH database under accession numbers GSE147082 and GSE118828. The datasets were processed using Seurat and integrated with Harmony. A keratin signature (KRT1, KRT2, KRT9, KRT10) was computed using Seurat’s AddModuleScore function and visualized in UMAP space.

For cell-cell communication analysis, the mean expression of IFI44L was calculated within the Monocyte/Macrophage population for each sample. Receptor genes were retrieved from the OmnipathR package and filtered based on a minimum expression level of log2(3) in T cells. Subsequently, mean receptor expression was calculated for each sample. Pearson correlation analysis was performed between IFI44L expression in Monocytes/Macrophages and receptor expression in T cells, with Benjamini-Hochberg correction applied to account for multiple testing.

### Differential Expression and Gene Set Enrichment Analysis

Differentially expressed proteins were identified using the limma R package, with patient identity and tumor site included as a covariate. GSEA was conducted using the fgsea package in R. The Gene Ontology gene sets from MSigDB were used. Pre-ranked gene lists, generated based on t-statistics (for infiltrated vs. non-infiltrated tumors) or Pearson correlation coefficients (for co-expressed genes), served as input. An FDR threshold of 0.05 was applied.

### The External Transcriptomic and Multicolor Immunofluorescence Dataset

The processed transcriptomics and immunofluorescence data was accessed at https://github.com/cansysbio/HGSOC_TME_Heterogeneity. The number of CD4+, CD8+ and Treg counts were averaged across each region in a sample and converted to a fraction of total counts for downstream analyses. The relationship between IFI44L expression and the mean expression of KRT1, KRT2, KRT9, and KRT10 with T cell infiltration was determined using independent linear regression models. The resulting coefficients were compared using a T-test.

### The External Proteogenomic Dataset

The processed proteomics data was accessed at https://pdc.cancer.gov/pdc/cptac-pancancer, and the TCGA molecular subtype annotations were available from the supplemental information of the paper. The mean protein expression of KRT1, KRT2, KRT9, and KRT10 was calculated in each sample, and compared between the TCGA molecular subtypes using an Anova test.

### T and B Cell Clonal Composition Analysis

The raw Nanostring counts data were accessed under accession number EGAS00001002839 and normalized using DESeq2. The genes identified as upregulated in infiltrated samples (logFC > 0; p value < 0.05) and non-infiltrated samples (logFC < 0; p value < 0.05) were scored using the GSVA package and compared between the immune phenotypes characterized by Zhang et al. The metrics defining T cell clonal composition data were accessed from the supplementary data.^1^ The partial Spearman correlation was calculated between the signatures of infiltrated and non-infiltrated samples, and metrics quantifying T cell clonal composition, controlling for the patient of origin.

### Ethics approval and consent to participate

Full ethical approval was obtained before sample collection from patients (Mater Misericordiae University Hospital Ethical Board—ref. ref 1/378/1954) and healthy volunteers (University College Dublin Ethical Board—ref. LS-21-06-Silva-Brennan). Informed consent was obtained for all individuals included in this study.

## Availability of data and material

The data that support the findings of this study have been deposited on MassIVE under ID MSV000094907 and are publicly available as of the date of publication. The principal analysis code used to analyze data and generate the results presented here are released through Zenodo. Any additional information required to reanalyze the data reported in this paper is available from the lead contact upon request.

## Competing Interests

Daniel C. Liebler is a stockholder and employee of Inotiv, Inc., which provides analytical services described. Salisha Hill and Ryan D. Morrison are employees of Inotiv. The remaining authors declare no competing interests.

## Funding

This publication has emanated from research conducted with the financial support of Precision Oncology Ireland, a Consortium of 5 Irish Universities, 6 Irish Charities, and 7 Industry Partners, which is funded by the Science Foundation Ireland Strategic Partnership Programme, under Grant number [18/SPP/3522]. Funding was also received from the Ireland East Hospital Group and the National Maternity Hospital Foundation. Biobanking activities at UCD-GOG are supported by the UCD Clinical Research Centre.

## Authors’ Contributions

Conceptualization, D.B., D.E., and K.G.; methodology, D.E., K.G., A.T., A.F., J.McC., S.H., R.M., D.L.; formal analysis, D.E.; investigation, D.E., K.G.; resources, D.B., D.L.; data curation, K.G., D.L.; writing – original draft, D.E.; writing – review & editing, D.B., W.K., D.E.; visualization, D.E.; supervision, D.B., W.K.; project administration, D.B.

---

**Supplementary Figure 1.**
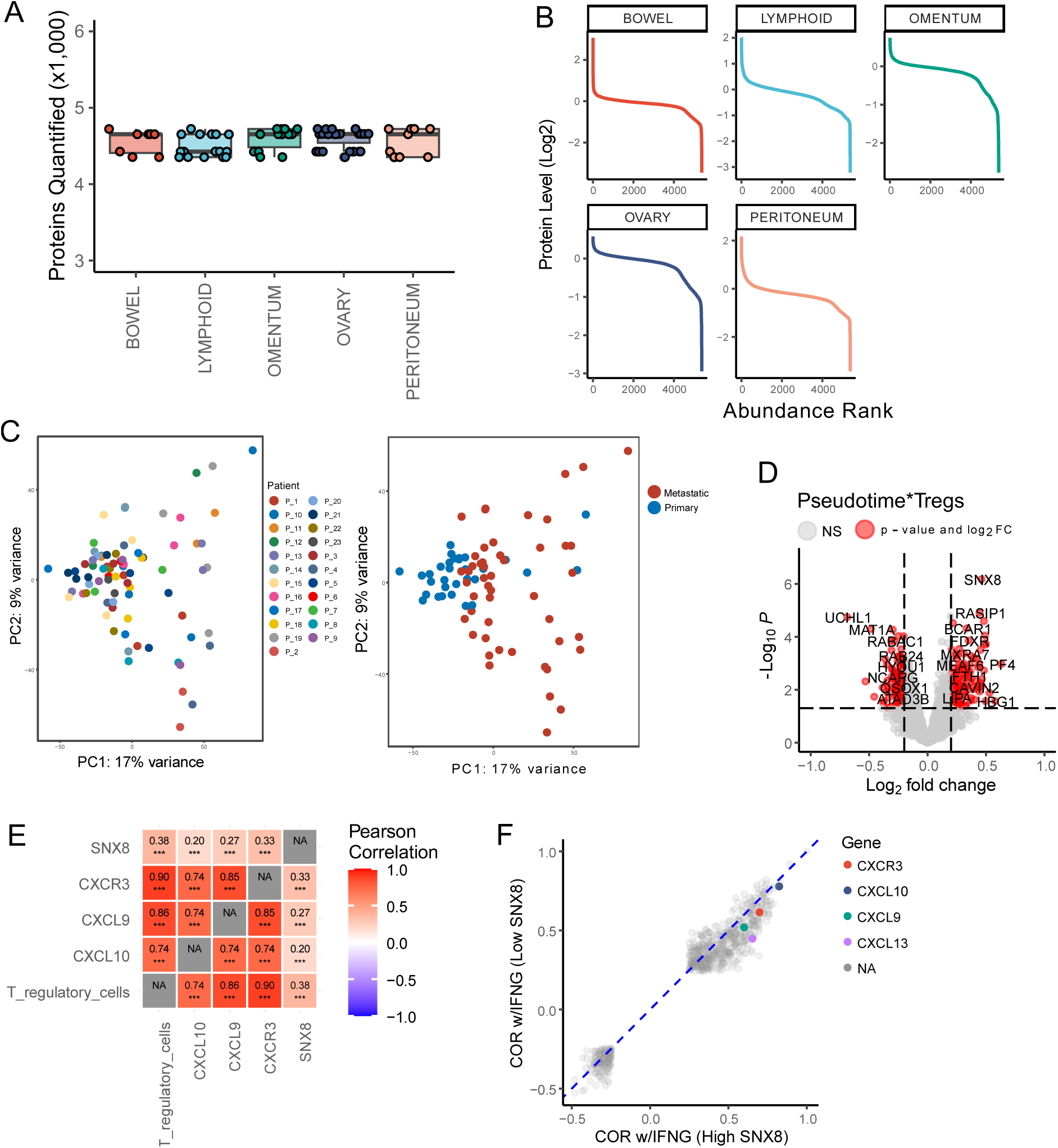
**A)** The number of proteins identified in each tumor site. **B)** The protein expression level and corresponding rank for each tumor site (dynamic range). **C)** Principal component analysis on the proteomic profiles of HGSOC tumor samples, annotated by patient identity and tumor site, respectively. **D)** Proteins with increasing (logFC > 0) or decreasing (logFC < 0) correlations with Tregs as a function of pseudotime. **E)** The correlation of Treg abundance with SNX8, CXCL9/10 and CXCR3 in ovarian cancer samples from TCGA. The color scale and numbers represent the correlation coefficients (*** = p < 0.01). **F)** The correlation between gene expression and IFNγ signaling in TCGA samples with either high or low expression of SNX8 (median cut-off). Genes to the right of the diagonal line have a stronger correlation with IFNγ signaling in samples with high SNX8 expression.

**Supplementary Figure 2.**
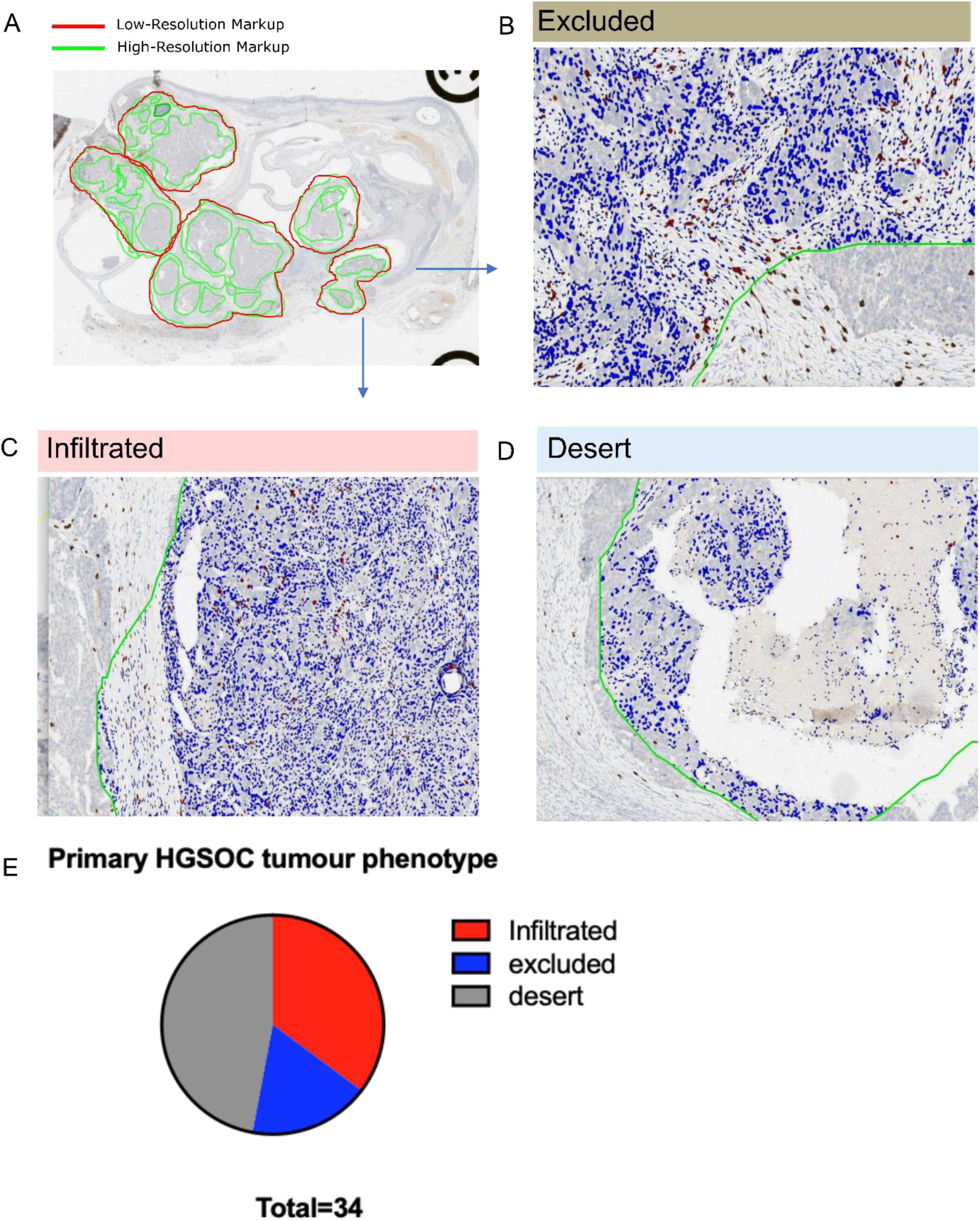
**A)** A primary ovarian sample stained for CD8+ T cells. The red lines denote the approach with fewer analyzed regions, while the green lines indicate the increased number of regions analyzed. **B-D)** x20 magnifications of the same sample demonstrating regions with an excluded, infiltrated, and desert immune phenotype, respectively. **E)** The proportion of each immune phenotype in the primary ovarian sample. 34 regions were examined at x20 magnification.

**Supplementary Figure 3.**
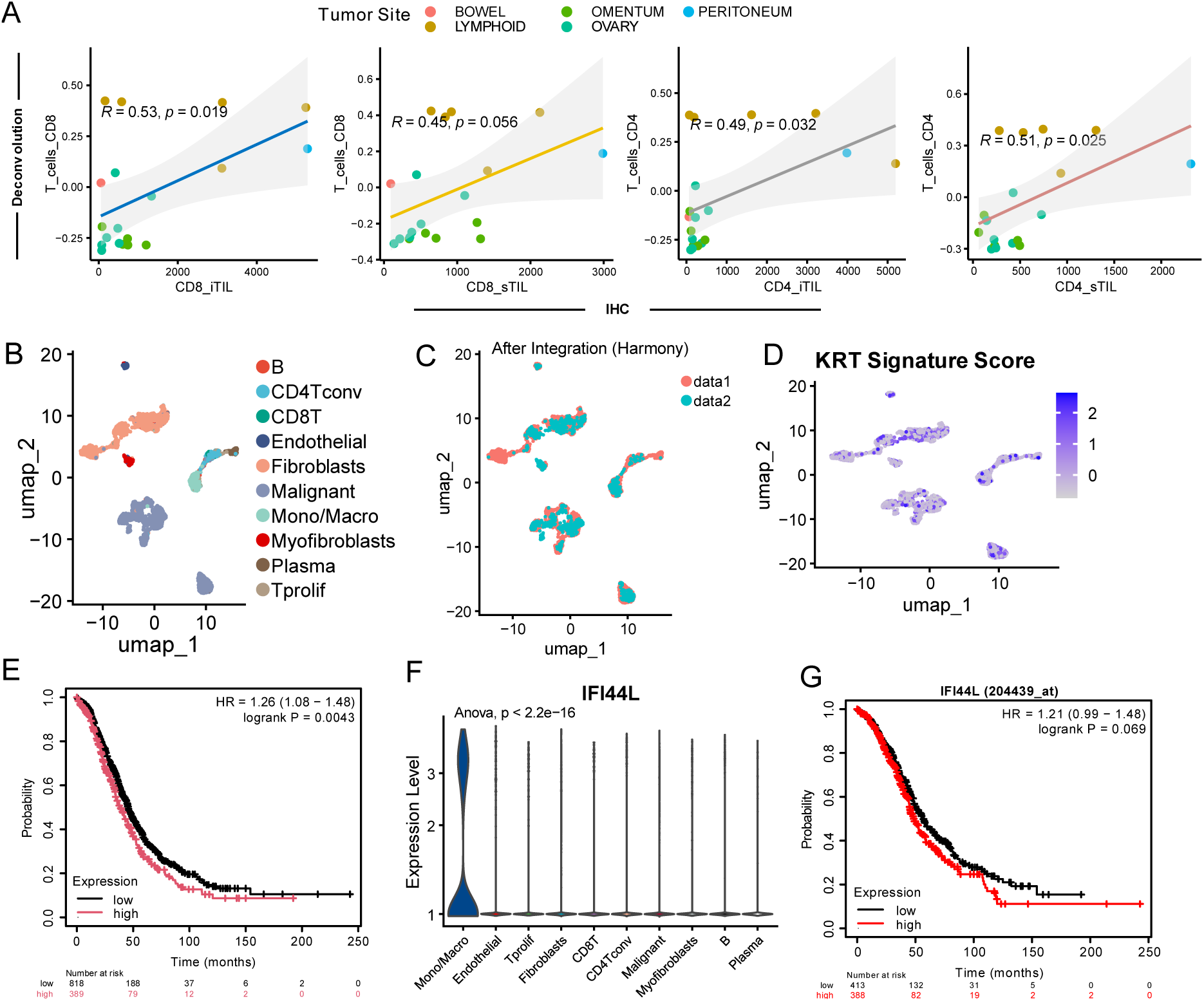
**A)** The correlation between TIL densities from IHC and deconvolution derived estimates of T cell abundances from global proteomics, calculated separately for CD8+ T cells (two left panels) and CD4+ T cells (two right panels). **B-D)** UMAP of cells from two integrated single cell RNA-seq datasets for ovarian cancer, annotated according to cell type, dataset of origin, and keratin signature score, respectively. The keratin signature contains KRT1, 2, 9 and 10. **E)** Kaplan Meier overall survival curve for high and low expression of keratin (KRT1, 2, 9 and 10). **F)** The expression of IFI44L in each cell type from the integrated single cell RNA-seq datasets. **G)** Kaplan Meier overall survival curve for high and low expression of IFI44L. Datasets from TCGA and GEO were used for survival analysis, available from.^23^

## REFERENCES

1. Zhang, A.W., McPherson, A., Milne, K., Kroeger, D.R., Hamilton, P.T., Miranda, A., Funnell, T., Little, N., Souza, C.P.E. de, Laan, S., et al. (2018). Interfaces of Malignant and Immunologic Clonal Dynamics in Ovarian Cancer. Cell 173, 1755–1769.e22. 10.1016/j.cell.2018.03.073.

2. Jiménez-Sánchez, A., Memon, D., Pourpe, S., Veeraraghavan, H., Li, Y., Vargas, H.A., Gill, M.B., Park, K.J., Zivanovic, O., Konner, J., et al. (2017). Heterogeneous Tumor-Immune Microenvironments among Differentially Growing Metastases in an Ovarian Cancer Patient. Cell 170, 927–938.e20. 10.1016/j.cell.2017.07.025.

3. Zhang, L., Conejo-Garcia, J.R., Katsaros, D., Gimotty, P.A., Massobrio, M., Regnani, G., Makrigiannakis, A., Gray, H., Schlienger, K., Liebman, M.N., et al. (2003). Intratumoral T Cells, Recurrence, and Survival in Epithelial Ovarian Cancer. New England Journal of Medicine 348, 203–213. 10.1056/NEJMoa020177.

4. Nabhan, M., Egan, D., Kreileder, M., Zhernovkov, V., Timosenko, E., Slidel, T., Dovedi, S., Glennon, K., Brennan, D., and Kolch, W. (2023). Deciphering the tumour immune microenvironment cell by cell. Immuno-Oncology and Technology 18, 100383. 10.1016/j.iotech.2023.100383.

5. Campbell, K.R., and Yau, C. (2018). Uncovering pseudotemporal trajectories with covariates from single cell and bulk expression data. Nat Commun 9, 2442. 10.1038/s41467-018-04696-6.

6. Eckert, M.A., Coscia, F., Chryplewicz, A., Chang, J.W., Hernandez, K.M., Pan, S., Tienda, S.M., Nahotko, D.A., Li, G., Blaženović, I., et al. (2019). Proteomics reveals NNMT as a master metabolic regulator of cancer-associated fibroblasts. Nature 569, 723–728. 10.1038/s41586-019-1173-8.

7. Hennessy, B.T., Coleman, R.L., and Markman, M. (2009). Ovarian cancer. The Lancet 374, 1371–1382. 10.1016/S0140-6736(09)61338-6.

8. Nieman, K.M., Kenny, H.A., Penicka, C.V., Ladanyi, A., Buell-Gutbrod, R., Zillhardt, M.R., Romero, I.L., Carey, M.S., Mills, G.B., Hotamisligil, G.S., et al. (2011). Adipocytes promote ovarian cancer metastasis and provide energy for rapid tumor growth. Nat Med 17, 1498–1503. 10.1038/nm.2492.

9. Coward, J., Kulbe, H., Chakravarty, P., Leader, D., Vassileva, V., Leinster, D.A., Thompson, R., Schioppa, T., Nemeth, J., Vermeulen, J., et al. (2011). Interleukin-6 as a Therapeutic Target in Human Ovarian Cancer. Clin Cancer Res 17, 6083–6096. 10.1158/1078-0432.CCR-11-0945.

10. Jiménez-Sánchez, A., Cast, O., and Miller, M.L. (2019). Comprehensive Benchmarking and Integration of Tumor Microenvironment Cell Estimation Methods. Cancer Research 79, 6238–6246. 10.1158/0008-5472.CAN-18-3560.

11. Wei, J., Guo, W., Lian, H., Yang, Q., Lin, H., Li, S., and Shu, H.-B. (2017). SNX8 mediates IFNγ-triggered noncanonical signaling pathway and host defense against Listeria monocytogenes. Proceedings of the National Academy of Sciences 114, 13000–13005. 10.1073/pnas.1713462114.

12. Chen, B.-J., Zhao, J.-W., Zhang, D.-H., Zheng, A.-H., and Wu, G.-Q. (2022). Immunotherapy of Cancer by Targeting Regulatory T cells. International Immunopharmacology 104, 108469. 10.1016/j.intimp.2021.108469.

13. Desbois, M., Udyavar, A.R., Ryner, L., Kozlowski, C., Guan, Y., Dürrbaum, M., Lu, S., Fortin, J.-P., Koeppen, H., Ziai, J., et al. (2020). Integrated digital pathology and transcriptome analysis identifies molecular mediators of T-cell exclusion in ovarian cancer. Nature Communications 11, 5583. 10.1038/s41467-020-19408-2.

14. Sade-Feldman, M., Yizhak, K., Bjorgaard, S.L., Ray, J.P., de Boer, C.G., Jenkins, R.W., Lieb, D.J., Chen, J.H., Frederick, D.T., Barzily-Rokni, M., et al. (2018). Defining T Cell States Associated with Response to Checkpoint Immunotherapy in Melanoma. Cell 175, 998–1013.e20. 10.1016/j.cell.2018.10.038.

15. Jiménez-Sánchez, A., Cybulska, P., Mager, K.L., Koplev, S., Cast, O., Couturier, D.-L., Memon, D., Selenica, P., Nikolovski, I., Mazaheri, Y., et al. (2020). Unraveling tumor– immune heterogeneity in advanced ovarian cancer uncovers immunogenic effect of chemotherapy. Nature Genetics 52, 582–593. 10.1038/s41588-020-0630-5.

16. Lu, S.X., De Neef, E., Thomas, J.D., Sabio, E., Rousseau, B., Gigoux, M., Knorr, D.A., Greenbaum, B., Elhanati, Y., Hogg, S.J., et al. (2021). Pharmacologic modulation of RNA splicing enhances anti-tumor immunity. Cell 184, 4032–4047.e31. 10.1016/j.cell.2021.05.038.

17. Korsunsky, I., Millard, N., Fan, J., Slowikowski, K., Zhang, F., Wei, K., Baglaenko, Y., Brenner, M., Loh, P., and Raychaudhuri, S. (2019). Fast, sensitive and accurate integration of single-cell data with Harmony. Nat Methods 16, 1289–1296. 10.1038/s41592-019-0619-0.

18. Olalekan, S., Xie, B., Back, R., Eckart, H., and Basu, A. (2021). Characterizing the tumor microenvironment of metastatic ovarian cancer by single-cell transcriptomics. Cell Reports 35, 109165. 10.1016/j.celrep.2021.109165.

19. Shih, A.J., Menzin, A., Whyte, J., Lovecchio, J., Liew, A., Khalili, H., Bhuiya, T., Gregersen, P.K., and Lee, A.T. (2018). Identification of grade and origin specific cell populations in serous epithelial ovarian cancer by single cell RNA-seq. PLOS ONE 13, e0206785. 10.1371/journal.pone.0206785.

20. Baptista, A.P., Roozendaal, R., Reijmers, R.M., Koning, J.J., Unger, W.W., Greuter, M., Keuning, E.D., Molenaar, R., Goverse, G., Sneeboer, M.M.S., et al. (2014). Lymph node stromal cells constrain immunity via MHC class II self-antigen presentation. eLife 3, e04433. 10.7554/eLife.04433.

21. Petralia, F., Ma, W., Yaron, T.M., Caruso, F.P., Tignor, N., Wang, J.M., Charytonowicz, D., Johnson, J.L., Huntsman, E.M., Marino, G.B., et al. (2024). Pan-cancer proteogenomics characterization of tumor immunity. Cell 187, 1255–1277.e27. 10.1016/j.cell.2024.01.027.

22. Thorsson, V., Gibbs, D.L., Brown, S.D., Wolf, D., Bortone, D.S., Ou Yang, T.-H., Porta-Pardo, E., Gao, G.F., Plaisier, C.L., Eddy, J.A., et al. (2018). The Immune Landscape of Cancer. Immunity 48, 812–830.e14. 10.1016/j.immuni.2018.03.023.

23. Győrffy, B. (2024). Integrated analysis of public datasets for the discovery and validation of survival-associated genes in solid tumors. Innovation 5. 10.1016/j.xinn.2024.100625.

24. Jiang, H., Tsang, L., Wang, H., and Liu, C. (2021). IFI44L as a Forward Regulator Enhancing Host Antituberculosis Responses. Journal of Immunology Research 2021, 5599408. 10.1155/2021/5599408.

25. DeDiego, M.L., Martinez-Sobrido, L., and Topham, D.J. (2019). Novel Functions of IFI44L as a Feedback Regulator of Host Antiviral Responses. Journal of Virology 93, 10.1128/jvi.01159-19. 10.1128/jvi.01159-19.

26. Türei, D., Korcsmáros, T., and Saez-Rodriguez, J. (2016). OmniPath: guidelines and gateway for literature-curated signaling pathway resources. Nat Methods 13, 966–967. 10.1038/nmeth.4077.

27. Dunlock, V.E. (2020). Tetraspanin CD53: an overlooked regulator of immune cell function. Med Microbiol Immunol 209, 545–552. 10.1007/s00430-020-00677-z.

28. Shevach, E.M. (2009). Mechanisms of Foxp3+ T Regulatory Cell-Mediated Suppression. Immunity 30, 636–645. 10.1016/j.immuni.2009.04.010.

29. Gotot, J., Gottschalk, C., Leopold, S., Knolle, P.A., Yagita, H., Kurts, C., and Ludwig-Portugall, I. (2012). Regulatory T cells use programmed death 1 ligands to directly suppress autoreactive B cells in vivo. Proceedings of the National Academy of Sciences 109, 10468–10473. 10.1073/pnas.1201131109.

30. Anadon, C.M., Yu, X., Hänggi, K., Biswas, S., Chaurio, R.A., Martin, A., Payne, K.K., Mandal, G., Innamarato, P., Harro, C.M., et al. (2022). Ovarian cancer immunogenicity is governed by a narrow subset of progenitor tissue-resident memory T cells. Cancer Cell 40, 545–557.e13. 10.1016/j.ccell.2022.03.008.

31. Goode, E.L., Block, M.S., Kalli, K.R., Vierkant, R.A., Chen, W., Fogarty, Z.C., Gentry-Maharaj, A., Tołoczko, A., Hein, A., Bouligny, A.L., et al. (2017). Dose-Response Relationship of CD8+ Tumor Infiltrating Lymphocytes and Survival Time in High-Grade Serous Ovarian Cancer. JAMA Oncol 3, e173290. 10.1001/jamaoncol.2017.3290.

32. Elpek, K.G., Lacelle, C., Singh, N.P., Yolcu, E.S., and Shirwan, H. (2007). CD4+CD25+ T Regulatory Cells Dominate Multiple Immune Evasion Mechanisms in Early but Not Late Phases of Tumor Development in a B Cell Lymphoma Model1. The Journal of Immunology 178, 6840–6848. 10.4049/jimmunol.178.11.6840.

33. Shimizu, J., Yamazaki, S., and Sakaguchi, S. (1999). Induction of Tumor Immunity by Removing CD25+CD4+ T Cells: A Common Basis Between Tumor Immunity and Autoimmunity. The Journal of Immunology 163, 5211–5218. 10.4049/jimmunol.163.10.5211.

34. Egan, D., Kreileder, M., Nabhan, M., Iglesias-Martinez, L.F., Dovedi, S.J., Valge-Archer, V., Grover, A., Wilkinson, R.W., Slidel, T., Bendtsen, C., et al. (2023). Small Gene Networks Delineate Immune Cell States and Characterize Immunotherapy Response in Melanoma. Cancer Immunology Research 11, 1125–1136. 10.1158/2326-6066.CIR-22-0563.

35. Curiel, T.J., Coukos, G., Zou, L., Alvarez, X., Cheng, P., Mottram, P., Evdemon-Hogan, M., Conejo-Garcia, J.R., Zhang, L., Burow, M., et al. (2004). Specific recruitment of regulatory T cells in ovarian carcinoma fosters immune privilege and predicts reduced survival. Nature Medicine 10, 942–949. 10.1038/nm1093.

36. Harmon, C., Zaborowski, A., Moore, H., St. Louis, P., Slattery, K., Duquette, D., Scanlan, J., Kane, H., Kunkemoeller, B., McIntyre, C.L., et al. (2023). γδ T cell dichotomy with opposing cytotoxic and wound healing functions in human solid tumors. Nat Cancer 4, 1122–1137. 10.1038/s43018-023-00589-w.

37. Ahmadzadeh, M., Pasetto, A., Jia, L., Deniger, D.C., Stevanović, S., Robbins, P.F., and Rosenberg, S.A. (2019). Tumor-infiltrating human CD4+ regulatory T cells display a distinct TCR repertoire and exhibit tumor and neoantigen reactivity. Science Immunology 4, eaao4310. 10.1126/sciimmunol.aao4310.

38. Norell, H., Carlsten, M., Ohlum, T., Malmberg, K.-J., Masucci, G., Schedvins, K., Altermann, W., Handke, D., Atkins, D., Seliger, B., et al. (2006). Frequent Loss of HLA-A2 Expression in Metastasizing Ovarian Carcinomas Associated with Genomic Haplotype Loss and HLA-A2-Restricted HER-2/neu-Specific Immunity. Cancer Research 66, 6387–6394. 10.1158/0008-5472.CAN-06-0029.

39. Yost, K.E., Satpathy, A.T., Wells, D.K., Qi, Y., Wang, C., Kageyama, R., McNamara, K.L., Granja, J.M., Sarin, K.Y., Brown, R.A., et al. (2019). Clonal replacement of tumor-specific T cells following PD-1 blockade. Nature Medicine 25, 1251–1259. 10.1038/s41591-019-0522-3.

40. Wu, T.D., Madireddi, S., de Almeida, P.E., Banchereau, R., Chen, Y.-J.J., Chitre, A.S., Chiang, E.Y., Iftikhar, H., O’Gorman, W.E., Au-Yeung, A., et al. (2020). Peripheral T cell expansion predicts tumour infiltration and clinical response. Nature 579, 274–278. 10.1038/s41586-020-2056-8.

41. Hu, Z., Artibani, M., Alsaadi, A., Wietek, N., Morotti, M., Shi, T., Zhong, Z., Santana Gonzalez, L., El-Sahhar, S., Carrami, E.M., et al. (2020). The Repertoire of Serous Ovarian Cancer Non-genetic Heterogeneity Revealed by Single-Cell Sequencing of Normal Fallopian Tube Epithelial Cells. Cancer Cell 37, 226–242.e7. 10.1016/j.ccell.2020.01.003.

